# *When feeling is better than seeing*: Adult Zebrafish Ignore Wide-Field Optic-Flow in Laminar, but not Turbulent Hydrodynamic Environments

**DOI:** 10.64898/2026.03.30.715425

**Authors:** S. Dave, J.C. Liao

## Abstract

Many animals navigate their world largely by seeing and feeling it. To disentangle these visual and mechanosensory contributions, we developed a virtual reality assay targeting the optomotor response in adult wild-type zebrafish swimming against flow. By projecting dynamic visual patterns onto the walls of a variable-speed flow tank, we decoupled wide-field optic flow from hydrodynamic velocity. We then tested fish responses to abrupt visual perturbations while they held station in the unsteady wake behind a bluff body. These perturbations reliably elicited compensatory optomotor responses, with fish aligning to the direction of the moving stimulus. Notably, this behavior was absent in uniform flows, suggesting that fish prioritize visual input when predictive lateral line signaling is compromised. We propose that this sensory shift serves to optimize swimming energetics in turbulent wakes. Extending this framework, we further show that zebrafish swimming against flow, whether alone or in groups, exhibit heightened escape responses to looming visual stimuli. Together, our findings reveal that fish sensory strategies are not fixed but dynamically tuned to hydrodynamic context: favoring visual cues in turbulent environments and lateral line input in uniform flows.

**Graphical Abstract:** 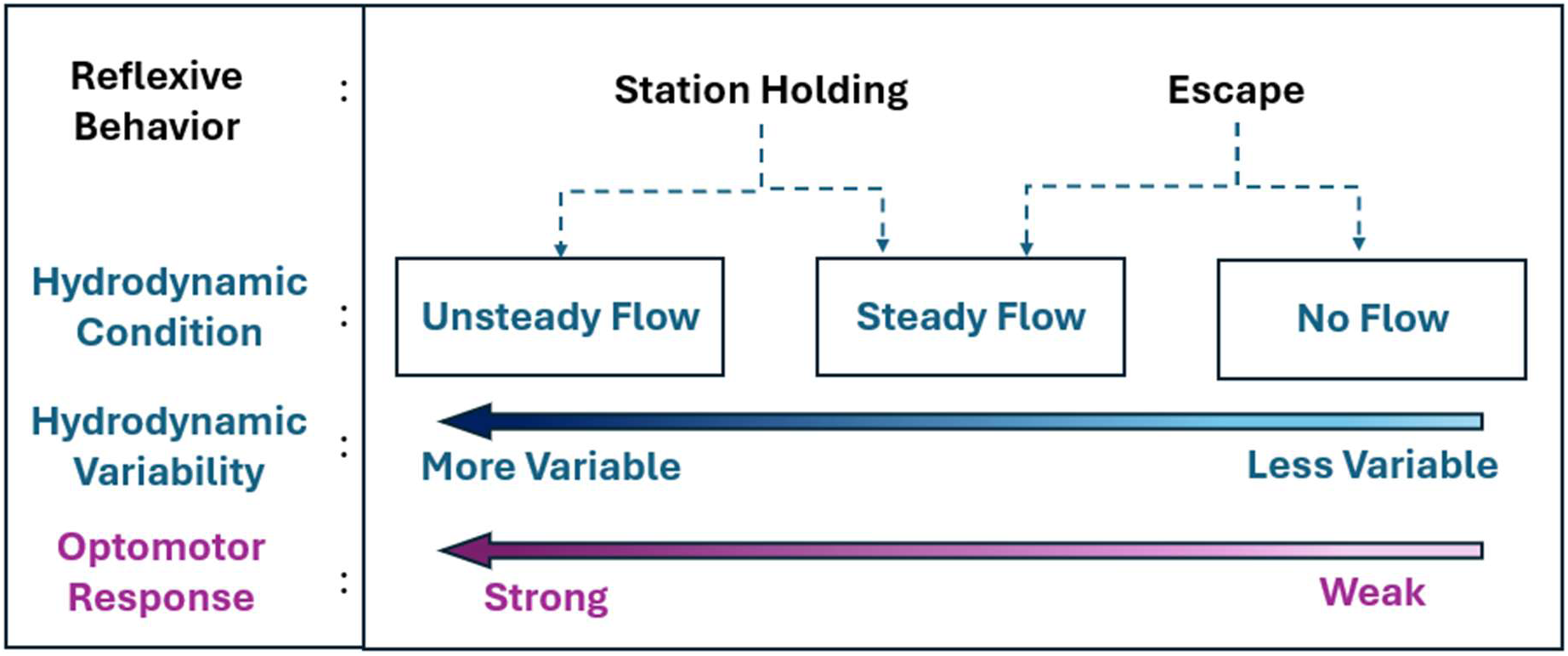

## Introduction

Sensory cues from different modalities often arrive simultaneously or overlap in sequence, providing animals with a rich and redundant source of information to navigate novel and complex environments. A growing acknowledgment of the integration and conflict dynamics of multiple sensory modalities (vision is coupled with olfaction, air flow, vestibular, etc.) has led to new insights in interpreting and understanding behavioral responses at the organismal level.

Animals moving through water experience vastly different challenges from terrestrial animals. In particular, the higher density and viscosity of water, the forces currents can create, and the relative lack of light characterize aquatic and marine habitats where vertebrate life originated. For over 400 million years, fishes have evolved to integrate mechanosensory and visual information to shape their behavior and ecology. The lateral line and visual system, detecting water flow and light, play a critical role in this integration, enabling fish to navigate and respond to their environment. The lateral line is an ancient mechanosensory system that predates the evolution of visual systems in vertebrates, and detects predators, prey, conspecifics and water currents. Specifically, genes associated with mechanosensation appear earlier in evolution than those related to visual processing (Šestak et al., 2013). This suggests that sensing water flow and pressure was a foundational capability for early aquatic vertebrates. However, despite the growing interest in understanding how multiple sensory systems orchestrate behavior (Dallmann et al., 2023; Sharma & Sponberg, 2023), our understanding of mechanosensation and vision in aquatic animals remains static and poorly understood.

In contrast to the highly stochastic nature of ambient turbulent flows, which, when interacting with environmental structures (rocks, vegetation, other animals), produce vortices of widely varying spatial and temporal scales, the boundary layer flow over a swimming fish’s skin (and thus lateral line system) is relatively predictable and repeatable (Gray, 1968). When flow interacts with a simple geometric shape like a cylinder, a vortex street can be generated. Fish can recapture the energy of these vortices to hold station in flow (e.g. maintain position relative to the Earth frame of reference). Fish holding station behind a cylinder show drastically reduced oxygen demands, saving up to half the cost of swimming compared to when swimming in laminar flow (Taguchi & Liao, 2011). Due to the turbulent regime in a vortex street, flow unpredictability causes destabilizing movements in these surfing fish (Liao et al., 2003b; Tritico & Cotel, 2010). Consistent with this, fish with an ablated lateral line avoid turbulence vortex streets and prefer to station holding in areas of smoother flow, suggesting a reliance on detecting flows to maintain position while station holding (Liao, 2006). Turbulence may limit the capabilities of the lateral line to signal normal swimming movements, where muscle commands align with sensory expectations. Predictable flow across the body is critical for the lateral line to generate an image of efficient swimming, enabling a proprioceptive function (Skandalis et al., 2021).

How does the hydrodynamic environment influence the reliance on visual information for station holding in adult zebrafish? We hypothesize that fish swimming in turbulence shift their sensory reliance from the lateral line towards vision. Here, we develop a novel virtual reality assay in a flow tank that decouples visual and hydrodynamic sensory inputs for freely swimming fishes. We use this approach to investigate the interplay between vision and the lateral line during optomotor and loom behaviors across laminar and turbulent flow conditions. We address previously inaccessible questions on the effect of wide-field visual on fish swimming and escape behaviors in flow. By decoupling and placing into conflict visual and hydrodynamic stimuli, our approach allows investigation into multi-agent and multisensory integration of fish behavior.

## Materials and Methods

### Animals

We used adult (>60 dpf) WT zebrafish, *Danio rerio* (body-length, mean± SE = 38.6 ± 0.7 mm, n=20 fish) raised in the UC Santa Barbara zebrafish facility and transferred to the experimental room >2 weeks prior to experiments. There, fish were maintained in two 10 L freshwater tanks maintained at 23 ± 0.5 °C) with a commercial aquarium heater (Eheim Co.), kept on a 12:12 light:dark cycle and fed commercial pellets *ad libitum* daily. Prior to the start of an experimental trial, an individual fish was introduced into the flow tank and left for 10 minutes to acclimatize at a current velocity of 10 cm s^-1^ (e.g. ∼2.5 body-lengths s ^-1^). We found this flow velocity would elicit the most reliable station holding response. All experimental trials were conducted in the afternoon (12:00-17:00 PST) in a room enclosed by blackout curtains. After data were collected from swimming trials, fish were euthanized with an over-dose of MS-222.

### Experimental Setup

#### Flow tank

All experiments were conducted using a custom-built 5 L recirculating flow tank with a working section of 22 x 7 x 7 cm (length x width x depth). Water flow was generated by a variable speed AC-DC series motor (Dayton model 2MO37A, 115v 1.5 Amp, Lake Forest Illinois USA) driving a propeller that circulated water through 2 sets of honeycomb collimators (1/8" aperture diameter) to generate uniform flow. Flow velocity was set at 10cm/s and verified by tracking suspended plastic particles within the working section of the flow tank. The cross-sectional area of the fish was less than 5% of the cross-sectional area of the flow tank, minimizing any solid blocking effects (Bell & Terhune, 1970). Water was filtered, aerated, and maintained at room temperature of 22.1 °C (SE=0.1°C) throughout the experiment. (See Figure-1A)

**Figure 1.**
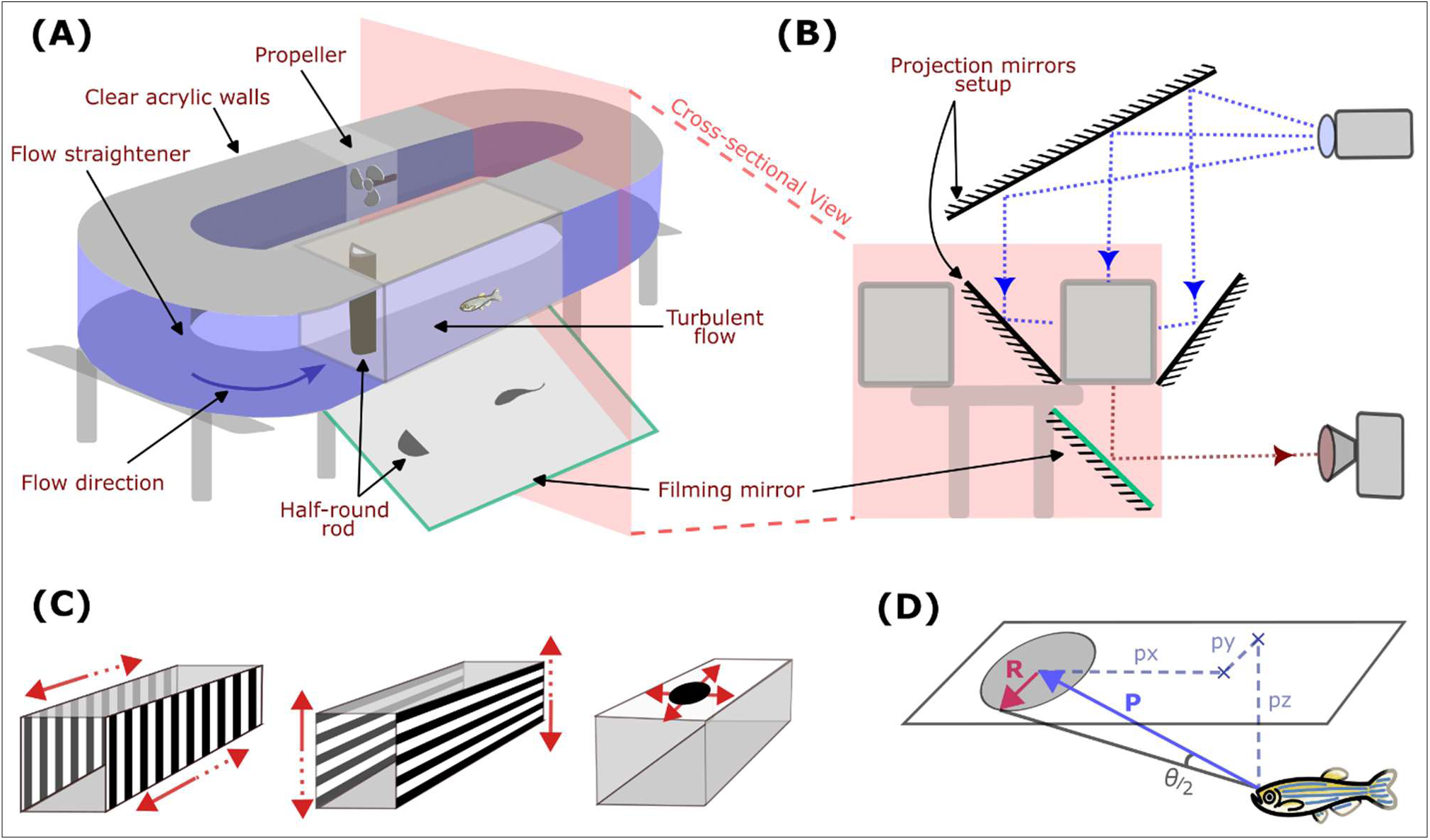
Schematic of experimental setup with the fish swimming flow-tank and visual stimulus projection. (A) A variable-speed flow tank placed on a stand with a filming-mirror placed underneath tilted at 45° angle to film the ventral view of the swimming fish. (B) Cross-sectional view of flow tank with two side-mirrors and a top-mirror used for visual stimulus projection, and filming mirror schematic. (C) Schematic of optical stimuli and their motion direction, where the patterns move simultaneously in the direction of sold arrows, and then for a separate treatment, move simultaneously in dashed-arrows directions. Vertical gratings for “optical push/pull” and horizontal gratings for “optical roll” treatments and the expanding loom stimulus on the top wall for escape trials. (D) Perceived threshold stimulus angle (θ) is calculated based on the 3D position of the escaping fish and the instantaneous radius of the loom stimulus.

#### Visual Projection

We used a system of mirrors (See Figure-1B) to project visual stimuli on customized rear-projection screens mounted to the flow tank walls. For wide-field grating patterns on the sidewalls, we varied orientation and motion (vertical gratings moving downstream/upstream; horizontal gratings moving up/down) and set the optic-flow speed as *ca.* 10 cm s^-1^ (spatial frequency=1.8 cm cycle^-1^, temporal frequency as 5.5 cycle s^-1^) to match hydrodynamic flow velocity. We recorded these moving projection patterns using a high-speed camera at 1000 frames s^-1^ to verify optic flow rates. For visual looming stimulus, we used an exponentially expanding (doubling-time: diameter=50ms, area=25ms) dark-circle with a bright background presented on the top-wall. Fish were acclimatized for 5 minutes before each trial with either horizontal or vertical gratings on sidewalls in case of station-holding, or plain white background on the top-wall in case of escape trials. For each fish trial, the treatments order was randomized to evoke a reflexive response and avoid contribution due to learning and memory.

#### Filming and Digitization

A monochromatic Chronos 1.4-HD high-speed video camera (1024 x 1024, 1000 frames/s, Burnaby BC Canada) was aimed at a front-surface mirror angled at 45° and placed directly below the working section to film the swimming kinematics and position of zebrafish (Figure-1B). An LED panel with white lights and overlayed diffuser was used to optimize the image contrast of the fish. In loom experiments, this panel was replaced with a projection screen for the top-projected loom stimulus with against a uniform background illumination. We used a semi-supervised machine-learning based tool, DeepLabCut (Mathis et al., 2018), that allowed us to reliably track 4 points on fish (head-tip, tail-tip, right- and left-pectoral fin base). To create a training-set, we manually labelled these 4 points in a total of 800 frames, randomly picked from 40 videos, and trained a neural network model for ∼1 million iterations to minimize the tracking error. We subsequently used this model to track points on more than 400,000 frames from 80 video recordings. Tracked videos using DeepLabCut were visually verified and approved before further analysis (Mathis et al., 2018).

### Experimental Design and Data Analysis

We studied the effects of hydrodynamic and visual conditions on freely swimming adult zebrafish using two widely studied behaviors, Station-holding and Escape behavior. Below are the experiment specific details for each behavior.

#### (i) Station-Holding Behavior

##### Water-flow conditions – Steady / Unsteady

Many fish routinely swim against water current to navigate, a behavior known as rheotaxis. These fish may experience Steady (laminar) or Unsteady (turbulent) water-flow. However, station holding fish would naturally experience unsteady water-flow as they swim behind rocks or bluff bodies to get help from eddies and conserve energy. We studied sensory prioritization in the position-maintaining fish in a flow tank and challenged them with steady and unsteady water-flow conditions. Steady flow conditions were generated by a custom 3D printed honeycomb flow-straightener. D-section cylinders (0.5-1 cm diameter) were added in an upstream location in the working section to create a distinct flow refuge to generate suitable unsteady flow (Liao et al 2003b).

##### Linear perturbations – Optical-Pull / Push

We first tested the role of wide field visual perturbation on a station holding adult fish. As the fish is swimming against water-flow, we externally provided wide-field optic-flow by suddenly moving the surroundings in (i.e., vertically oriented visual gratings) forward (upstream) or backwards (downstream) directions. The fish typically experience optic-flow either during self-movements or when external factors (such as water-flow) move them. Since our treatments induced the optic-flow analogous to fish moved by external factors, we call them ‘optical-push’ and ‘optical-pull’, based on the direction of optic-flow generated. ‘Optical-push’ is the treatment of moving visual surroundings from back to front of the fish, mimicking the optic-flow direction if fish are pushed backwards externally (e.g., sudden increase of water-flow speeds). Similarly, optical-pull treatment will suddenly move wide-field visual patterns from front to back (i.e., in a downstream direction) mimicking a scenario when a station holding fish is pulled forward due to external factors (e.g., sudden decrease of water-flow speeds). The prefix “optical” here refers to the fact that it’s a purely visual perturbations, rather than changing waterflow speed to move the fish.

##### Rotational perturbations – Optical-Roll

We further checked the effects of a conflicting optic-flow on station holding fish, where the optic-flow presented along rotational axis whereas the water-flow remained linear (along the body-axis). We moved visual surroundings (i.e., horizontally oriented visual gratings) in opposite directions on each side wall at the same spatial and temporal rates as the ‘Optical-Push / Pull’ treatments (see Methods: Visual Projections). We call them ‘optical roll’ treatments that would mimic optic-flow generated to the fish if it is externally rotated in Roll-direction (around the longitudinal-axis). The rotational optic-flow is either in clockwise (CW: right-side moving down, left moving up) or counterclockwise (CCW: right-side moving up, left moving down) directions from the fish’s perspective.

##### Data Analysis

To examine the effect on station holding behavior, we compared 1 second of fish trajectory data before and after the stimulus and quantified changes in body position, swimming velocity, and tail beat. We excluded the first 500ms of data after the stimulus onset from this analysis. Swimming velocity was divided into longitudinal (along the direction of flow) and lateral (side-ways) directional components to better explain the behaviors observations. Changes in trajectory pattern were quantified using Spearman’s rank correlation (rho), which measures monotonic changes in the position (where a constant upstream movement = 1, and downstream = −1). We also tracked the distal end of tail and quantified its average cyclic movement amplitude and frequency over a given duration for each trial.

#### (ii) Escape Behavior

##### Experimental Treatments

We studied another naturalistic behavior that is known to involve vision, an escape behavior, to further compare how fish use vision in different hydrodynamic conditions. We presented an exponentially expanding (looming) stimuli (see Methods: Visual Projections) at the top wall of swimming chamber while fish were challenged to swim against water-flow (Flow) and when at rest (No-Flow). We studied fish’s response when alone in the chamber (Single) or when in a group of five individuals (Group). The looming stimulus generally originated towards the upstream side of the chamber, giving better visibility to the fish swimming against the flow. However, we also carried out a treatment where the position is shifted towards the downstream side of the chamber (Downstream Loom), to study any possible effects arising due to the stimulus position.

##### Data Analysis

We studied fish’s reflexive escape responses involving sudden bending of the body, known as C-Start, upon presenting the looming stimulus. From the video recordings, we find fish’s instantaneous position w.r.t. the stimulus origin in horizontal plane ([px,py] cm) and the instantaneous stimulus radius (R cm) at the time of an escape response (i.e., onset of C-start reflex). Vertical position of the fish (pz cm) is considered as the flow-tank height, as the fish were found swimming at the bottom of the tank. From the escaping fish’s three dimensional position (P=[px,py,pz] cm) and stimulus radius (R), we can then calculate the threshold stimulus angle (theta) from the fish’s perspective that elicited the escape response (using the formula: theta = 2*arctan( R / Norm(P) ). We call this angle (theta) the Perceived Loom Angle. We also measured the escape response delay as the time difference between beginning of the looming stimulus and the C-start.

## Results

### Prioritizing vision depends on hydrodynamic environment

We quantified fish swimming performance and their responses to wide-field visual perturbation (Optical-Pull and -Push treatments, see methods: Exp. Design) while swimming against steady and unsteady water currents in the flow-tank. For each treatment, we observed responses from 4 adult zebrafish and repeated 4 trials for everyone, making a total 16 unique trials per treatment. We used Wilcoxon signed-rank test to compare behavior responses between the pre-stimulus and post-stimulus values.

### Fish do not rely on optic-flow to maintain swimming position in steady flows

During trials with steady (laminar) flow, fish’s swimming velocities in the streamwise direction remain unchanged after the stimulus onset when compared to the pre-stimulus adaptation period for Optical-Pull (velocities mean ± s.e.m.: pre-stim = −0.2 ± 1.2 cm/s, post-stim = 0.2 ± 1.2 cm/s; p = 0.90, n.s.) and, also for Optical-Push treatments (velocities mean ± s.e.m. : pre-stim = −0.1 ± 1.0 cm/s, post-stim = −0.9 ± 1.1 cm/s; p = 0.74, n.s.). Conversely, presenting exactly the same visual stimulus but now with unsteady (turbulent) currents elicited stereotypical compensatory optomotor response (Figure-2 C,D). In unsteady flow, swimming velocities in streamwise-directions reduces for Optical-Pull (velocities mean ± s.e.m.: pre-stim = −0.2 ± 0.4 cm/s, post-stim = −4.2 ± 0.6 cm/s; p = 0.0004), whereas it increases for Optical-Push treatment (velocities mean ± s.e.m.: pre-stim = −0.3 ± 0.6 cm/s, post-stim = 2.3 ± 0.6 cm/s; p = 0.009). This behavior can be explained as positive optomotor response (OMR), since the fish’s post-stimulus displacement is in the same direction as the visual projections on sidewalls, for both Optical-Pull and Push perturbations in unsteady water currents. A compensatory positive OMR may help station-holding fish to maintain its position using visual feedback. Such OMR is missing for both the treatments in steady currents, suggesting a prominently non-visual sensory mechanism to maintain position.

**Figure 2.**
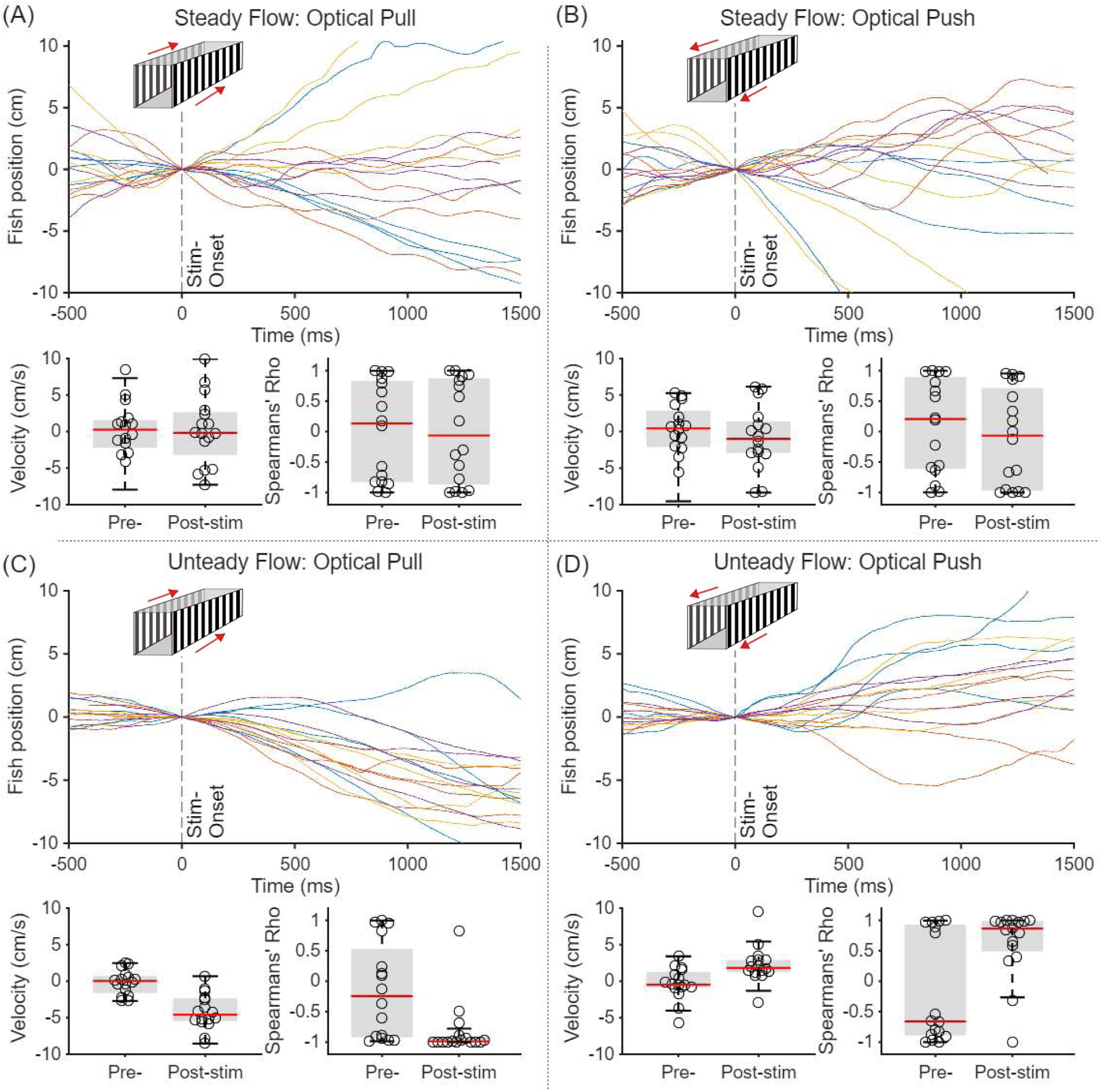
Quantifying station-holding fish’ response to wide-field optical perturbations in steady water-flow (A-B) and unsteady flow(C-D). (A,C) “Optical pull” stimulus entails visual patterns suddenly moving downstream at the stimulus-onset (vertical dashed-line). *Top-panel*: Fish’s longitudinal positions (along the streamwise direction) is plotted against time, where negative position values show downstream drift for “optical pull” (visual patterns moving downstream) and each color represents one individual fish (n=4 fish, 4 trials each); *Bottom-panel:* Comparing the change in fish’s swimming velocity and Spearman’s correlation co-efficient (representing degree of monotonous movements) before and after the stimulus onset. (B,D) Same plots as in part-A but for the “optical push” stimulus, where visual patterns move upstream at the onset (n=4 fish, 4 trials each).

### Fish move monotonously with wide-field optic flow in unsteady flows

We compared fish’s trajectories with a purely linear model trajectory (with a slope=1) and quantified Spearman’s correlation coefficient (rho) to study the monotonicity of their responses for each treatment. Rho values of 1 and −1 represent a perfectly monotonous change of position in the forward and in backwards direction, respectively. We found that fish swimming in unsteady flows showed a higher monotonous change in their body position post-stimulus onset. Rho value for Optical-Pull (rho ± s.e.m : pre-stim = −0.2±0.2, post-stim = −0.8±0.1, p=0.009) representing a monotonous downstream drift, and for Optical-Push (rho ± s.e.m : pre-stim = −0.2±0.2, post-stim = 0.6±0.1, p=0.04) representing largely monotonous upstream surge, post stimulus onset in unsteady flows. However, when in steady flows, the rho values are spread across the spectrum with averages close to zero representing a lack of overall monotonous movements (Optical-Pull: pre-stim = 0.0±0.2, post-stim = 0.0±0.2, p=0.99, n.s.; Optical-Push: pre-stim = 0.1±0.2, post-stim = −0.1±0.2, p=0.74) (Figure-2).

### Compensatory OMR only present for streamwise (longitudinal) direction

For both steady and unsteady hydrodynamic conditions, the post-stimulus change in position happened in the streamwise direction (Figure-2, longitudinal), in the same direction as the visual stimulus. Cross stream (e.g. lateral) position, velocity, and Spearman rho values remain unchanged post-stimulus (Figure-S1, all p>0.4, n.s.). We further found that when presenting optical-roll perturbations (i.e., clockwise and counter-clockwise), fish did not display an OMR either in longitudinal or lateral directions (Figure-S2, all p>0.4, n.s.). We also note that fish did not turn, change its swimming direction or show an escape response (e.g., C-start) to any of our wide-field visual stimuli. Raw trajectories and summary statistics values of the means, s.e.m and p-values are included in supplemental material (Supp. Figure-1,2, Supp.Table-S1).

Thus, our results of visual perturbation experiments during station holding suggest that adult zebrafish use optic-flow to maintain position while swimming against unsteady water streams, and the compensatory optomotor response to optic-flow depends on the direction of optic-flow and hydrodynamic conditions.

#### Fish sensitivity to visual threats alters with hydrodynamic conditions

As our previous experiment showed that fish swimming against water currents did not trigger escape responses (e.g. C-start) to wide-field visual perturbations, next we presented an exponentially expanding, purely visual looming stimulus on the dorsal wall of the flow tank and studied their behavioral responses. We quantified fish’s escape attempts responses (C-start) due to looming stimulus presented during swimming against water current (Flow) and no currents (No-Flow), both for the individual trails (Single) or when schooling (Group) (Figure-3). We used Wilcoxon rank-sum test to compare quantities between Flow and No-Flow treatments.

**Figure 3.**
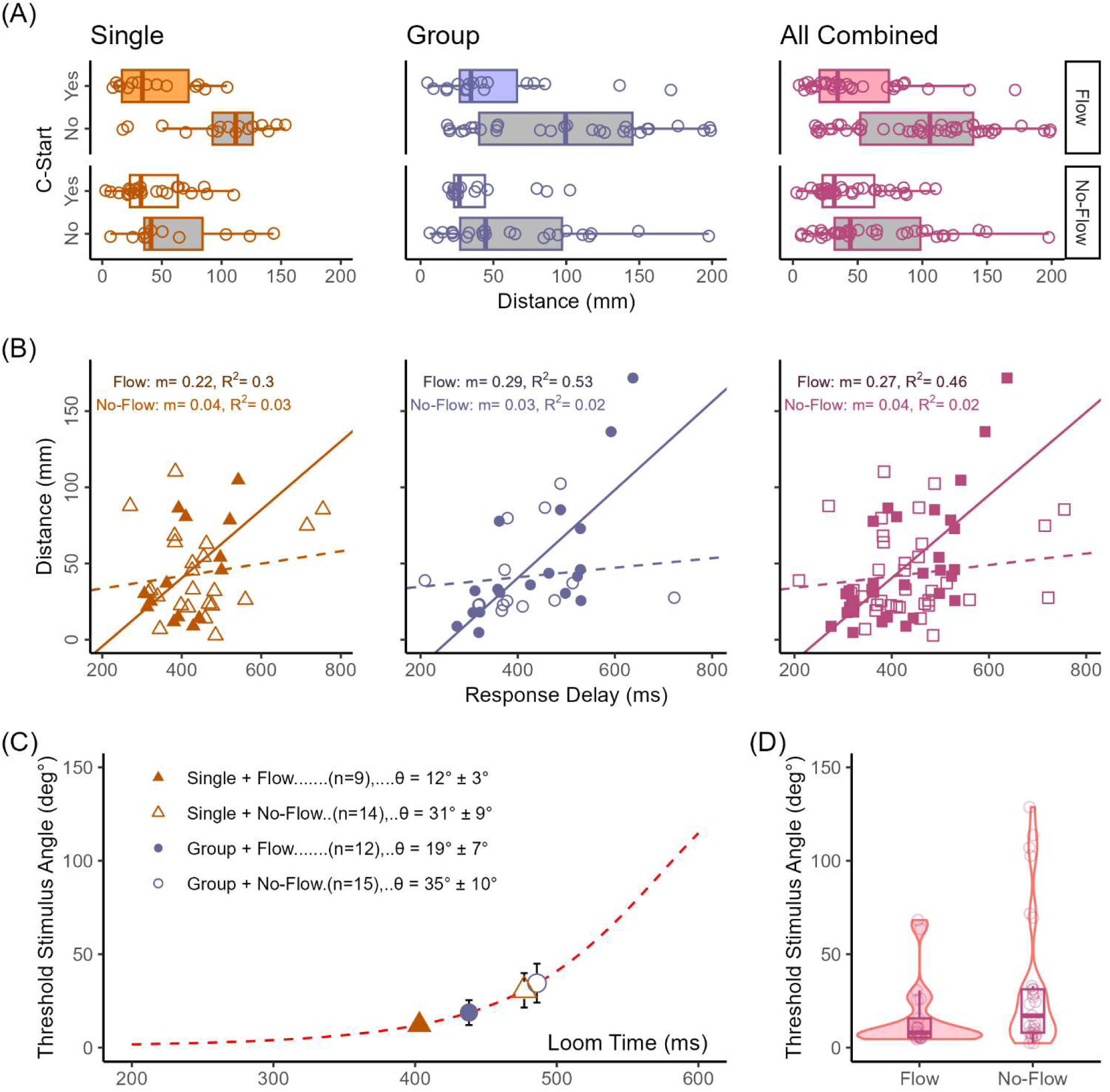
Comparison of escape response (C-start) due to a purely visual looming stimulus during “flow” (swimming against water current) and “no-flow” (still water) conditions in individual (“Single”) and school (“Group”) of fish. (A) Presence or absence of escape trigger (C-start) is classified for “flow” (top) and “no-flow” (bottom) conditions, and plotted against fish’s swimming distance from the stimulus center point. (B) Similar to part-A but now projecting the looming stimulus on a different part (i.e., towards the downstream end of the flow-tank). (C-D) For the escaping fish, their swimming distance and response delay are compared between “flow” and “no-flow” conditions. A higher slope indicates stronger correlation between those quantities. (E) Perceived angle of the threshold stimuli (θ), which is found using fish’s 3D position and instantaneous radius of looming stimulus, for all trials are plotted on a representative looming stimulus (Dashed line) that indicates the angular expansion of the looming stimulus over time. (F) The same threshold stimulus angles for all the fish are compared between “flow” and “no-flow” trials. A lower angle represents quicker escape.

### Escape Distance-Delay relationship depends on hydrodynamic conditions

Adult zebrafish escape responses for both Single and Group trials showed that the escaping fish were positioned closer to the stimulus origin as compared to the non-escaping fish. This relationship between the escape response presence and the distance from stimulus are consistent across hydrodynamic conditions, Flow (distance mean ± s.e.m.: Escape = 48.5±6.3 mm, n=36; No-Escape = 93.7±7.2 mm, n=53; p=9.1e-6) and No-Flow (Escape = 41.8±4.2 mm, n=41; No-Escape = 60.3±6.4 mm, n=48; p=0.002) (Figure-3A). Thus, we considered fish’s position when triggering a C-start response in our subsequent analysis of the behavior.

We also quantified the escape response delay since the start of looming stimulus expansion for each of the escaping fish in all trials and studied its relationship with fish’s distance from stimulus to find whether closely positioned fish escaped faster. Flow trials show a stronger positive relationship (slope m = 0.20 for Single, m = 0.24 for Group) when fishing swimming against current than in No-Flow trials (slope m = 0.05 in Single, m = 0.04 in Group) (Figure-3 B). We then performed linear regression analysis because raw slope values are susceptible to any mismatch in axis ranges for distance and delay. Combining escape events from Single and Group trials, we found a stronger escape distance-delay correlation for the Flow treatment (Pearson coefficient = 0.67, n=36, p<0.001) than in No-Flow conditions (Pearson coefficient = 0.21, n=41, n.s.). This analysis shows that the fish that were swimming against flow triggered C-start sooner when located closer and later when further away from the stimulus. Whereas the non-swimming fish in general did not show such response, suggesting they just responded to the presence of the threatening stimulus.

### Station holding fish show a lower threshold-angle to looming stimulus

The escape behavior may be better studied by quantifying the threshold stimulus angle for each escaping individual because it would account for both the response delay and fish’s distance from the stimulus. We calculated the threshold stimulus angle of the looming stimulus from the perspective of the fish in three-dimensional space at the time of escape to test whether station holding fish (Flow trials) responded to the angular expansion faster than in No-Flow trials, as it would be predicted by their slope-values (Figure-3B). To maintain consistency across each responses, we considered all trials for which the looming stimulus was originating within the lateral-visual field (153° field on right and left side each; n=50 fish) for the given fish positions, and excluded data with the stimuli in their blind-spot (21° field on rear side; n=5 trials) and binocular (33° field on front; n=22 trials) (Pita et al., 2015). This allowed us to better compare our data across Flow and No-Flow trials because fish’s body orientations are not always facing upstream when not swimming, as they would while swimming against water currents (No-Flow trials).

We found that station holding fish triggered an escape at a lower stimulus angle in Flow as compared to stationary fish in No-Flow (Figure-3C). These angles are plotted on a model looming stimulus to visualize their median threshold angles on the expanding visual stimulus in temporal axis. Finally, combining the threshold stimulus angles from each escaping fish across all the Single and Group trials, we found a lower threshold for fish swimming against water current (Flow: 15.9°±3.9° deg, n=21) than fish in stationary water (No-Flow: 32.6°±6.8° deg, n=29) (Figure-3D). A lower threshold angle (p=0.017) to trigger a C-start reflex suggest increased sensitivity to the visual threat stimuli.

Together, our results highlight the relationship between the optic-flow and hydrodynamic flow, where fish swimming in complex water currents elevate their response to optic-flow perturbations.

## Discussion

The OMR is crucial for an animal’s ability to maintain its position, and is particularly vital for fish navigating dynamic aquatic habitats. Our research offers new insights into the adaptive strategies of sensory modalities in fishes, revealing how they prioritize visual and mechanosensory cues based on the predictability of their hydrodynamic environment, and how these strategies influence other crucial behaviors like escape responses.

### Developmental and Hydrodynamic Influences on OMR

Our study of adult zebrafish reveals a distinct OMR compared to that observed in larvae. While larval zebrafish typically exhibit a positive OMR, swimming with moving bars to reduce optic flow (Olive et al., 2016), adults demonstrate a negative OMR, swimming against the visual motion (Bak-Coleman et al., 2015). This developmental shift likely reflects the maturation and calibration of sensory and motor systems (Kohashi et al., 2012). Larvae, with less developed sensory systems and limited proprioception, may primarily rely on visual cues for body displacement. In contrast, fully integrated adult systems allow for the development of robust expectations of flow and the accumulation of error-driven motor learning experiences (Montgomery et al., 2002; Skandalis et al., 2021). This calibration enables adults to form precise expectations of how lateral line (and likely vestibular) inputs correspond with visual inputs in predictable hydrodynamic environments like still water or steady flow.

However, this calibrated expectation for sensory information, typically associated with steady swimming kinematics, is violated in turbulent flows. We suggest that the positive OMR observed in larvae arises because they initially rely heavily on visual inputs as they gain the experience necessary to fully calibrate their mechanosensory and visual systems. A compelling avenue for future investigation would be to precisely determine the developmental stage at which this OMR switch occurs, and whether this transition can be accelerated or delayed by specific environmental conditions like light levels or turbulence.

Beyond neural development, the hydrodynamic regime itself imposes distinct challenges and opportunities for sensory processing. Our findings indicate that fish dynamically adjust their reliance on mechanosensation versus vision in a context-dependent manner, revealing a previously unrecognized prioritization strategy driven by hydrodynamic predictability, which correlates with Reynolds number (Re). Larval fish operate at low Reynolds numbers (Re < 100), where viscous forces dominate and flow patterns are highly predictable. In this regime, consistent lateral line reafference from self-motion likely allows larvae to prioritize visual inputs for OMR, as mechanosensory input provides a stable internal reference. Conversely, adult zebrafish operate at high Reynolds numbers (Re > 1000), routinely encountering turbulent flows characterized by complex, unpredictable vortices in natural aquatic environments. When uniform flow interacts with bluff bodies, the resulting vortices can significantly alter swimming kinematics (Liao et al., 2003a; Sutterlin & Waddy, 1975). Consequently, the lateral line system receives less predictable input than during self-generated swimming in uniform flow (Crapse & Sommer, 2008). While vortices can be detected (Chagnaud et al., 2007), their presence likely diminishes the lateral line’s capacity to provide a consistent signal for self-motion or external flow, substantially reducing its reliability as an accurate indicator of effective swimming in turbulence (Skandalis et al., 2021).

Similar principles of context-dependent sensory reweighting have been observed in terrestrial systems. For example, hawkmoths tracking moving flowers modulate their reliance on vision and mechanosensation depending on ambient luminance (Sharma & Sponberg, 2023). In bright light, visual gain increases while mechanosensory gain decreases; in dim light, the reverse occurs. This mirrors our findings in zebrafish, where lateral line input becomes less reliable in turbulent flow, prompting a shift toward visual cues. In both cases, animals preserve behavioral performance—flower tracking in moths, station-holding and escape in fish—by flexibly reconfiguring sensory strategies. These parallels suggest that adaptive sensory prioritization, rather than fixed integration, may be a widespread solution to environmental uncertainty.

### Energetic Imperatives and Adaptive Sensory Prioritization

The diminished reliability of lateral line input in turbulent flows, coupled with the energetic imperative of station-holding, profoundly influences how fish perceive and react to their environment. The drive to conserve energy strongly shapes fish behavioral choices, particularly in complex flow fields, and consequently their reliance on specific sensory cues.

Fish exploit the energetic benefits of station-holding within vortical wakes generated behind bluff bodies (Liao, 2004; Stewart et al., 2016; Taguchi & Liao, 2011). This strategy, termed Karman Gaiting, involves precise positioning to reduce the energetic demands of swimming (Liao et al., 2003a, 2003b). Fish actively return to these energetically favorable regions after displacement, demonstrating a strong motivation to maintain such positions. Optimal station-holding often occurs at a specific saddle point downstream from the obstacle, where flow conditions minimize energy expenditure (Taguchi & Liao, 2011; Zdravkovich, 1997). Even a small displacement can lead to ejection into the higher-cost freestream, with oxygen consumption twice as high (Taguchi & Liao, 2011). These substantial energetic costs provide a powerful incentive for fish to actively maintain station in a specific region of the wake; a benefit absent in the viscous regime occupied by larval fish.

This drive leads to a marked shift in sensory prioritization. In predictable, uniform laminar flows, where lateral line inputs are reliable, OMR is absent or diminished. This suggests lateral line prioritization due to its direct, rapid, and continuous feedback on the body from the water flow that vision alone cannot replicate with equivalent fidelity or speed. However, when station-holding fish encounter the unpredictable, noisy hydrodynamic cues of a turbulent wake, vision rises to a paramount role. This results in the emergence of a positive OMR, where fish swim in the direction of a moving visual stimulus. For a fish maintaining a fixed position in a turbulent flow-field, such a visual strategy is highly adaptive. Ignoring visual cues would result in costly downstream drift, or risk ejection into the high-cost freestream. Interestingly, stationary near-field visual cues (e.g., the cylinder) proved insufficient to maintain station-holding in turbulence, indicating that broad, wide-field visual stimuli are required for this behavior.

### Role of the Efferent System and Sensory Conflict

A switch to visual reliance in turbulent flows suggests an underlying neural mechanism that actively modulates sensory input. We propose that the efferent system of the lateral line plays a critical role in this dynamic reweighting of sensory information. The efferent system cancels out flow information generated by self-movement (e.g., corollary discharge, (Crapse & Sommer, 2008)), allowing zebrafish to prune afferent information correlated with swimming motions and the fluid environment (Skandalis et al., 2021). This comparison of expectation and sensing enables the lateral line to convey external proprioceptive abilities in simple flow environments.

Building on this, we hypothesize that the visual system is also calibrated into this proprioceptive architecture. Sensing the predictable flow across the body during anticipated undulations suppresses behavioral responses stemming from wide-field visual inputs; hence, fish in steady flows do not react to optical perturbations. When holding station in turbulent flow, the hydrodynamic cues necessary for body positioning during swimming diminish, prompting fish to switch to visual cues to guide their behavior and maintain position. In these conditions, fish in our study show compensatory responses to optical push and pull treatments.

Importantly, when station-holding fish follow downstream-drifting bars, they do not turn and swim downstream, nor do they cease swimming and drift passively while still facing upstream. Instead, they maintain an upstream orientation and swim slower, using a reduced tailbeat frequency and amplitude. Active swimming is necessary to activate an efferent motor copy, which also improves control. Swimming, through corollary discharge, could calibrate flow expectations along the body, allowing fish to recognize less turbulent (e.g., more predictable) flows. Such an awareness is harder to execute when swimming downstream, given the anterio-posterior sensitivity bias of neuromasts (Munz, 1985). Through the lateral line efferent/afferent system, fish are continuously in touch with their environment. Undulating the body allows fish to detect changes in turbulence, as exemplified when fish, visually prompted to leave their station-holding region, eventually encounter freestream flow outside the wake. This would be impossible if fish drifted downstream straight-bodied, without the efferent activity to gauge body awareness.

Our hypothesis that the efferent system gates visual input is further supported by our sensory-conflict experiments. Our optical-roll experiments reveal that visual input becomes less influential when lateral line input aligns with an expected environmental model. This suggests that continuous hydrodynamic input to the lateral line system overrides sudden visual inputs, causing fish to disregard cross-stream moving bars when flow is moving downstream (this study, (Bak-Coleman et al., 2015)). In virtual rolling scenarios, the absence of dorsoventral flows typically detected by the lateral line, coupled with semicircular canal inputs that do not align with the rolling movements conveyed by the visual world, further highlights this sensory prioritization. This raises a crucial question: if the prioritization of the lateral line over the visual system is a fundamental strategy for navigating predictable environments, do these principles of context-dependent sensory weighting extend to other fundamental, survival-critical behaviors? To answer this, we next investigated if this sensory flexibility also applies to rapid, ecologically vital behaviors like the escape response, a well-characterized behavior fish use to flee predators based on auditory, visual, or lateral line stimuli (Mirjany et al., 2011).

### Vision-Dependent Escape Responses in Flow and Group Dynamics

For over a century, the escape response has been a focal point of research, primarily in individual fish within simplified hydrodynamic environments. While most studies have been conducted in still water, fish in nature often form groups as an anti-predator response (Nadler et al., 2021; Krause & Ruxton, 2002; Magurran, 1990) and frequently inhabit current-swept environments. Less understood, however, is the effect of group dynamics on escape behavior, where individuals must process not only visual threats but also information from moving group members.

We observed that fish initiated escape responses more frequently when in closer proximity to a loom stimulus, regardless of whether they were individuals or in a group, and whether they were swimming in flow or still water (Figure-3 A, B). However, the delay in response showed a stronger correlation with stimulus distance when fish were swimming in flow compared to still water (Figure-3 C, D). This positive correlation between distance and response delay suggests that fish are reacting to the angular expansion of the stimulus rather than merely its presence. When accounting for both stimulus distance and response delay, fish in flow exhibited a lower angular threshold for triggering an escape response. This finding aligns with the hypothesis that fish are more sensitive to visual threats while swimming in flow than in still water.

Our results suggest that the lateral line may be sensitized to compensate for potential constraints in escape trajectories within flowing water. Unlike in still water, where escape speeds and energetic costs are relatively equal in all directions, upstream escape paths in flow likely result in slower bursts and higher energetic costs compared to downstream or cross-stream paths (Domenici & Hale, 2019; Kohashi et al., 2012). This implies that faster responses in flow are a prominent component of station-holding behavior. Indeed, faster flows have been shown to elicit faster performance phenotypes in other fish species in the wild (Nadler et al., 2018).

The heightened visual sensitivity of zebrafish in challenging hydrodynamic environments demonstrates their adaptive flexibility in sensory processing. However, this behavioral adaptation frequently occurs within a social context, as zebrafish intrinsically associated in groups. This raises further questions about how collective dynamics influence threat perception and response. We found that individuals within a group exhibited a higher angular threshold for escape compared to solitary individuals. This suggests that schooling favors robustness to a stimulus rather than increased sensitivity, a phenomenon also documented in wild fish (Fahimipour et al., 2023). Interestingly, while schooling exposes individuals to unpredictable hydrodynamic stimuli generated by conspecifics, this appears to have a desensitizing effect on vision, a stark contrast to the OMR where individuals become more sensitive to visual threats.

### Rethinking the Lateral Line: Beyond Simple Flow Sensing

To more precisely dissect the independent contributions of the lateral line and vestibular systems to sensing turbulent flow, future experiments should aim to create hydrodynamic conditions that generate unpredictable flow across the body while ensuring that vortex strength and size do not displace the body. This approach would allow for the investigation of unreliable lateral line information concurrent with predictable vestibular input, a distinction not achievable in the present study due to the absence of lateral line ablation experiments.

Traditional antibiotic or genetic ablations of the lateral line, by primarily targeting hair cells, selectively remove afferent (incoming) flow information while leaving the efferent system intact. This suggests that such ablations do not isolate the study of flow sensing alone but instead introduce a more complex scenario that likely involves the unmasking of other sensory modalities, such as body sensing, which complicate behavioral interpretations. Building on this, we predict that fish with a non-functional lateral line, when exposed to uniform flows, will exhibit behaviors analogous to those observed in turbulent flow conditions. In these situations, alterations in the visual wide-field may elicit behavioral responses that would typically be disregarded in a uniform flow environment (e.g., antibiotic studies, (Liao, 2006)). This intricate interplay among sensory modalities may represent a fundamental behavioral mechanism that prioritizes robust, rapid signals from the lateral line system, and may have been necessary for animals before advanced visual systems were establish (Šestak et al., 2013).

## Conclusion

In summary, we demonstrate that wide-field visual inputs do not alter the behavior of adult zebrafish swimming in uniform flows. We argue that during uniform flow conditions, where hydrodynamic stimuli can be anticipated and compared to an internal model of movement, fish prioritize flow inputs from the lateral line system and/or the vestibular system over wide-field visual stimuli. In contrast, fish holding station in turbulent flows alter their behavior in response to wide-field visual inputs. We reason that the lateral line can no longer reliably predict flow along the body in unsteady flows as it can during uniform swimming. Because these fish may be less certain of their swimming state based on lateral line mechanoreception, vision emerges to play a larger role in directing behavioral responses. The greater energetic consequence of forfeiting position when station-holding behind a bluff body makes it particularly significant that visual inputs are acted upon once lateral line inputs become less predictable. Furthermore, while schooling, fish experience unpredictable hydrodynamic stimuli created by other individuals, yet they exhibit a decreased sensitivity to looming stimuli. Our work supports the idea that the prioritization of sensory modalities, rather than simple integration, is specific to flow environments. The ability to dynamically prioritize sensory inputs underscores the adaptive capacity of fish in complex and challenging environments. These findings emerge from an approach that embraces more complex and naturalistic experiments, and have important implications for fields ranging from neuroscience, collective behavior, and robotic control.

## Contributions

Conceptualization: J.C.L., S.D. Methodology: J.C.L., S.D.; Software: S.D. Validation: J.C.L., S.D.; Formal analysis: S.D.; Investigation: J.C.L., S.D.; Resources: J.C.L; Data curation: J.C.L., S.D.; Writing - original draft: J.C.L., S.D.; Writing - review & editing: J.C.L., S.D.; Supervision: J.C.L.; Project administration: J.C.L; Funding acquisition: J.C.L.

## Acknowledgements

We would like to thank Matteo Adorisio for help with experiments and preliminary analysis, and Eileen Hamilton for fish care. All protocols were approved by the University of Santa Barbara Institutional Animal Care and Use Committee. This research was supported in part by NSF Grant No. PHY-1748958, NIH Grant No. R25GM067110, Gordon and Betty Moore Foundation Grant No. 2919.01, and the Kavli Foundation. This research was additionally supported by the National Science Foundation Grants IOS 1257150, 2321275, and 1856237; NSF MPS/PHY 2102891 and NSF ENG/CMMI 2345913 to J. Liao, and National Institute on Deafness and Other Communication Disorders Grant R56DC020321 to J. Liao. The authors declare no competing interests.

## Supplementary Materials

**Supp. Figure 1.**
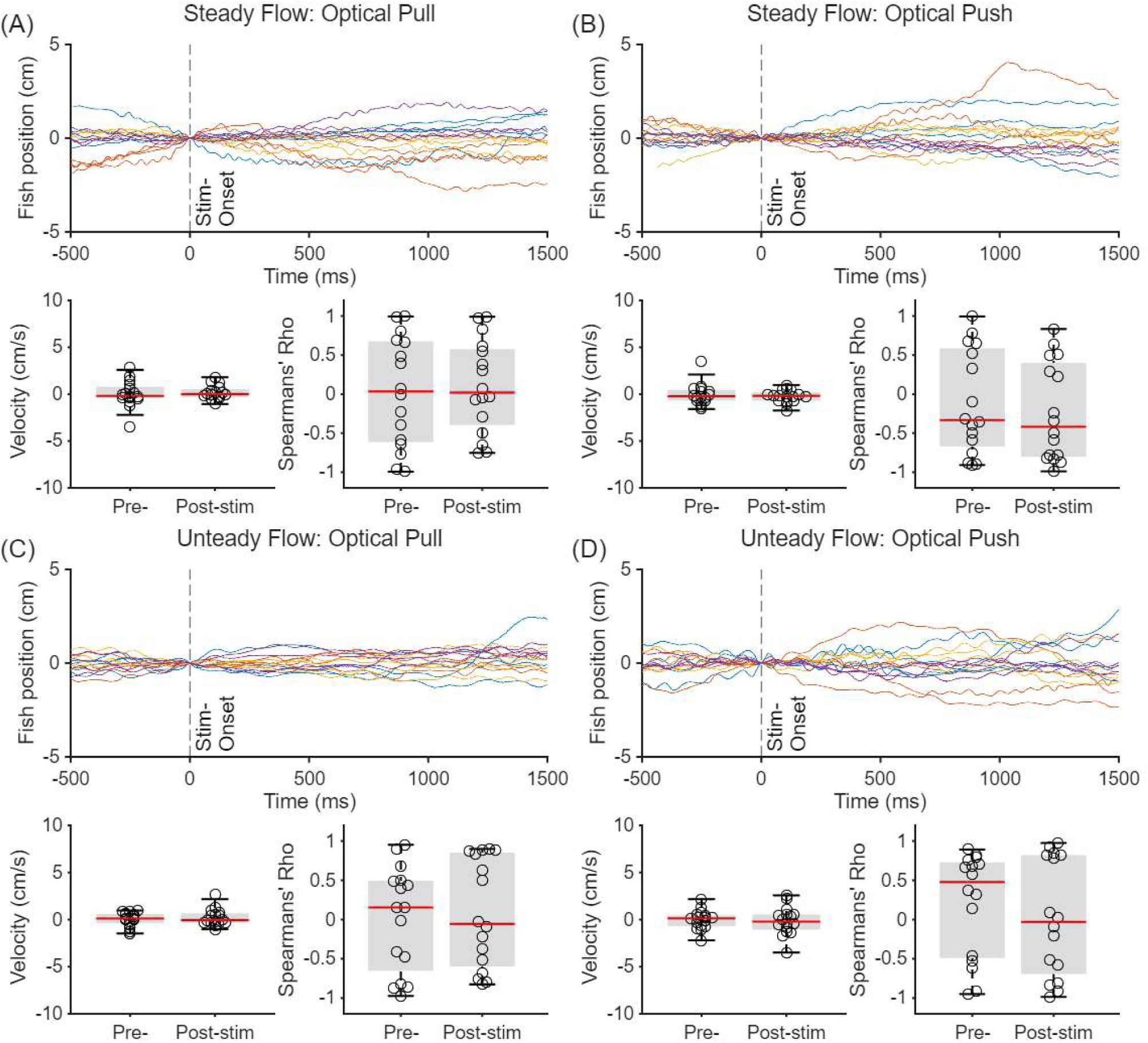
Raw trajectory data and velocity data for lateral movements in optical pull/push experiments for steady and unsteady flow (raw data for longitudinal movements, and plot legends are as in Fig-2).

**Supp. Figure 2.**
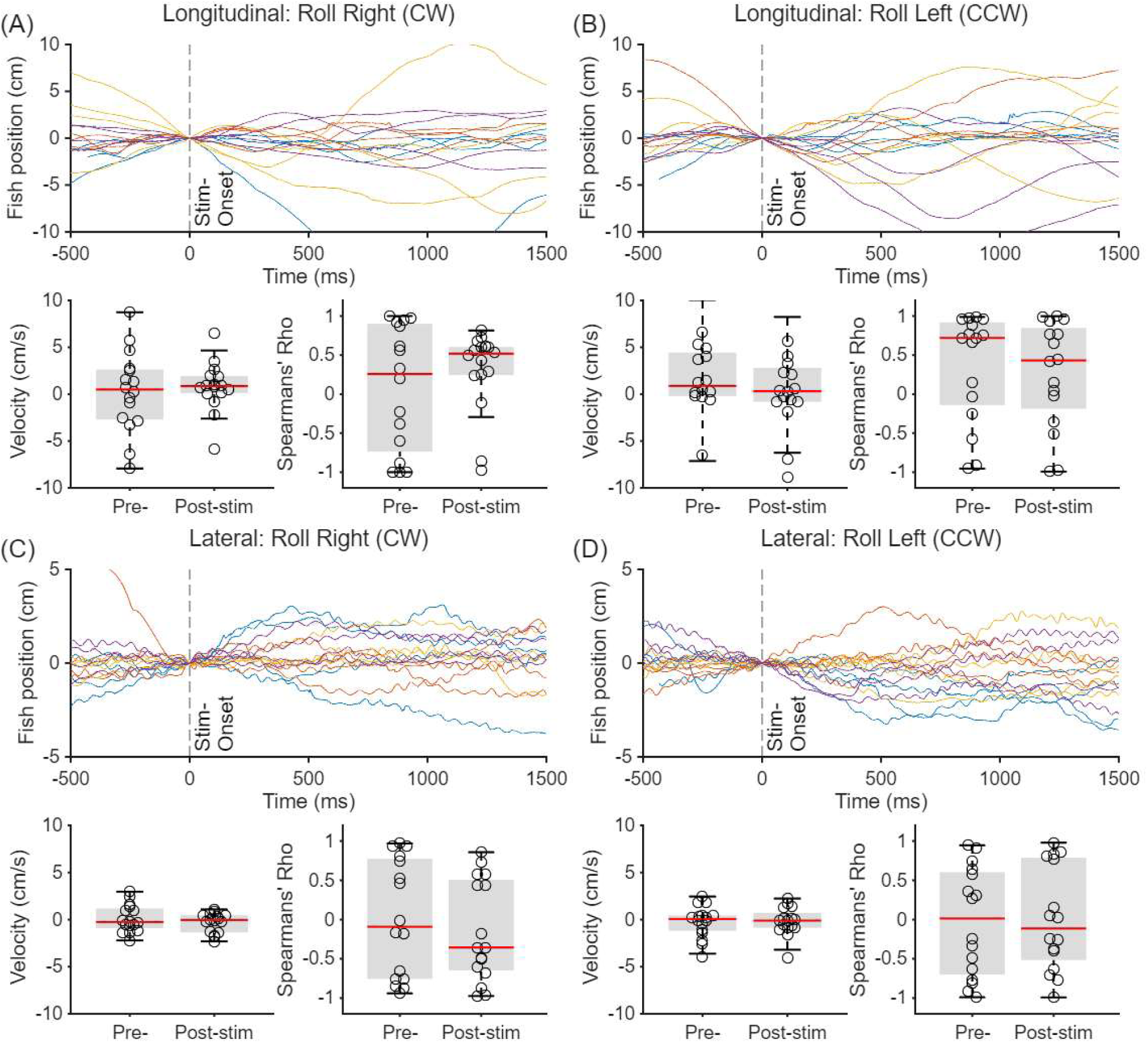
Raw data for optical roll perturbations (CW & CCW directions) for longitudinal and lateral direction movements. Plot legends are similar to explained in Fig-2 (A-B))

**Supp. Figure 3.**
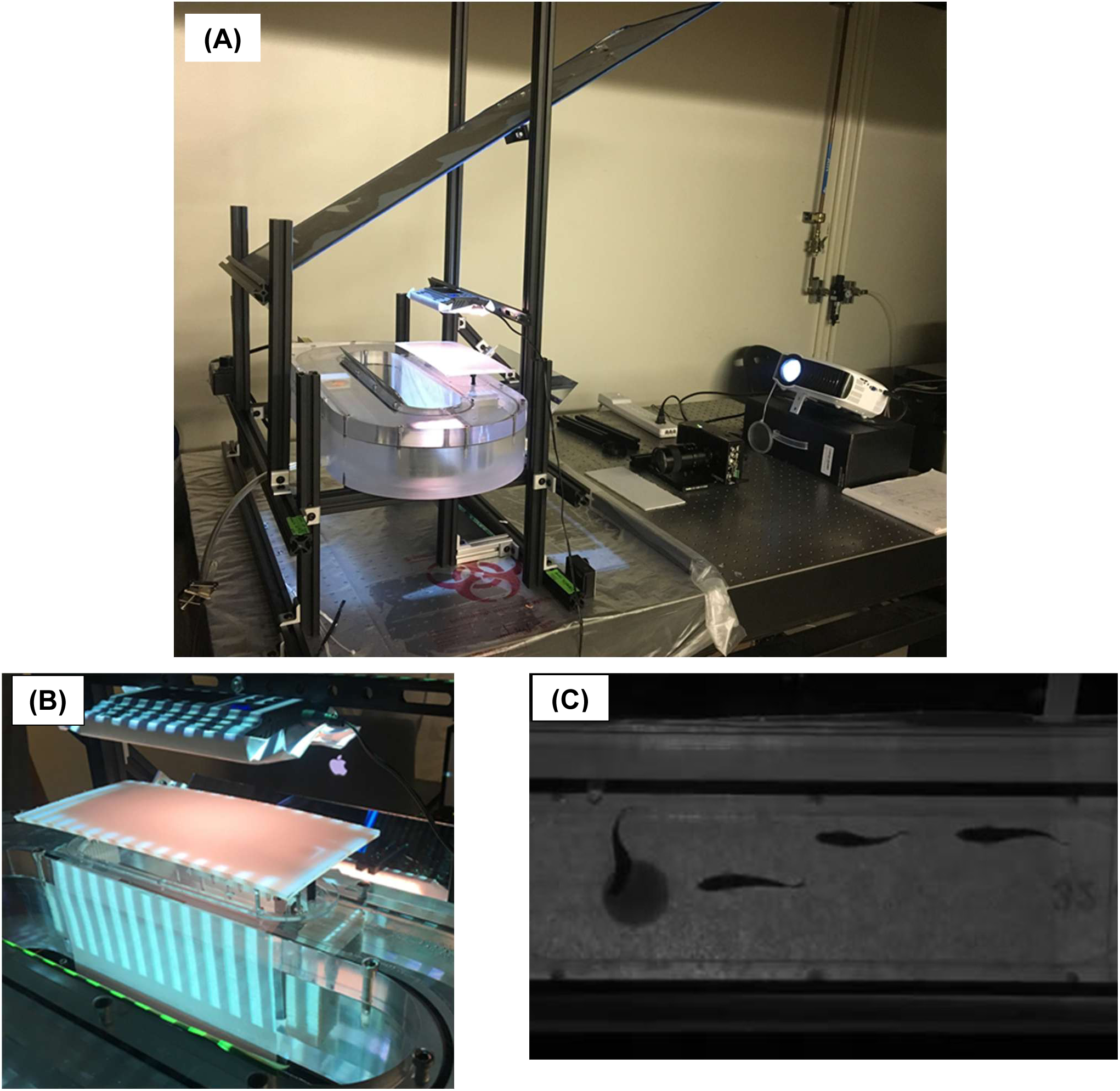
Images of the experiment setup. (A) Flow tank, Projection and Recording setup. (B) Example visual stimulus projected on a side-wall of swimming chamber. (C) A section of an image acquired by the camera of a group of fish swimming with an expanding looming stimulus in the background and the first fish (from upstream side) triggering a C-start reflex.

**Table S1:** Statistical summary of lateral-only motion in response to visual perturbation.

| Lateral-Only | Side Velocity mean ± s.e.m (cm/s) | | P-val | Side rho mean ± s.e.m | | P-val |
| --- | --- | --- | --- | --- | --- | --- |
|  | Pre-Stimulus | Post-Stimulus | Pre vs Post | Pre-Stimulus | Post-Stimulus | Pre vs Post |
| Optical-Pull (Steady) | 4.2E-04 ± 3.5E-03 | 1.5E-03 ± 2.0E-03 | 0.98 (n.s.) | 3.2E-02 ± 1.8E-01 | 1.0E-01 ± 1.5E-01 | 0.82 (n.s.) |
| Optical-Push (Steady) | -3.7E-04 ± 2.9E-03 | -2.2E-03 ± 1.7E-03 | 0.63 (n.s.) | -1.1E-01 ± 1.7E-01 | -2.3E-01 ± 1.6E-01 | 0.60 (n.s.) |
| Optical-Pull (Unsteady) | 3.6E-04 ± 1.8E-03 | 2.3E-03 ± 2.3E-03 | 0.67 (n.s.) | 1.5E-02 ± 1.7E-01 | 8.0E-02 ± 1.7E-01 | 0.86 (n.s.) |
| Optical-Push (Unsteady) | -1.6E-04 ± 2.5E-03 | -1.6E-03 ± 3.8E-03 | 0.74 (n.s.) | 2.0E-01 ± 1.7E-01 | 2.1E-02 ± 1.9E-01 | 0.46 (n.s.) |
| Optical-Roll (cw) | -2.5E-03 ± 4.2E-03 | -1.8E-03 ± 3.7E-03 | 0.98 (n.s.) | -2.6E-02 ± 1.7E-01 | 6.0E-03 ± 1.7E-01 | 0.90 (n.s.) |
| Optical-Roll (ccw) | 1.2E-03 ± 3.6E-03 | -3.3E-03 ± 2.6E-03 | 0.74 (n.s.) | 1.4E-02 ± 1.9E-01 | -1.5E-01 ± 1.6E-01 | 0.86 (n.s.) |

